# HPeV-3 predominated among *Parechovirus A* positive infants in the summer of 2013-2014 in Queensland, Australia

**DOI:** 10.1101/182436

**Authors:** Donna McNeale, Claire Y.T. Wang, Katherine E. Arden, Ian M. Mackay

## Abstract

Parechoviruses (HPeV) are not new viruses and are found in the respiratory tract and central nervous system of children and adults in conjunction with a range of acute illnesses. During an Australian outbreak of HPeV in the summer of 2013, we performed PCR-based screening and genotyping to determine whether ill Queensland infants were infected by HPeV. HPeVs were detected among 25/62 samples, identified as HPeV-3 from 23 that could be genotyped. These variants closely matched those occurring during and after the 2013 HPeV season. The inclusion of HPeV screening should be considered among acutely ill young infants during summer.

**Highlights:**

- HPeV-3 was the most common *Parechovirus A* genotype in Queensland summer of 2013/14
- HPeV testing should be routine among testing of infants with acute CNS - related symptoms
- HPeV is a seasonal virus
- Subgenomic phylogenetic analysis of HPeVs can be confounded by the presence of recombination

## Introduction

Members of the species *Parechovirus A* includes human parechovirus (HPeV) type 1 and 2. They were first identified in 1956, initially called echovirus 22 and 23 and in 2004, HPeV-3 was added.[1-4] There are currently 19 HPeV genotypes assigned to the genus *Parechovirus,* family *Picornaviridae*.[1] Infection has been associated with respiratory [5, 6] and gastrointestinal disease [6, 7] in adults. HPeV-3 in particular has been implicated as the principal cause of neonatal sepsis and meningitis and has been described as the most prevalent picornavirus in central nervous system (CNS)-related infections, sometimes with rash and seizure.[8-10] Young children are more likely than adults to suffer aseptic meningitis [6], encephalitis, flaccid paralysis and severe neonatal sepsis.[6, 11, 12] Long term neurodevelopmental sequelae from HPeV infection have also been reported.[13] Infections usually occur in warmer months with HPeV-1 and -3 the most often identified genotypes.[8, 14-18] Seroprevalence studies in Japan and Finland have noted >85% of adults have preexisting antibodies to HPeV-1 or 3.[3, 19] HPeVs are more resistant to alcohol-based hand disinfectant than influenza A virus and transmission is thought to be by droplets and after contact with contaminated surfaces.[20, 21] HPeV RNA can be shed in the faeces of infants for up to six months.[22]

In routine clinical practice, the primary focus remains to exclude treatable causes of CNS infection, including bacterial and herpesvirus infections.[10] The nature of modem screening technologies allows a range of other pathogens to also be sought during testing. Detection of HPeVs in stool or CSF samples is clinically useful for acute gastrointestinal and CNS disease, usually achieved using reverse transcriptase real-time polymerase chain reaction (RT-rtPCR) methods.[10, 23] Rapid and sensitive genotyping allows differentiation of previously assigned genotypes and can be achieved by characterising highly variable sequences spanning the junction of the viral protein (VP) coding regions VP3/VPl.[24]

During 2013, reports appeared which described HPeV-3 cases in young children in the Australian east coast state of New South Wales (NSW).[13, 18, 25, 26] These reports led us to offer PCR-based testing to seek out and characterize the genotype of *Parechovirus A* suspected to be causing illness in Queensland.

## Material And Methods

### Specimens

For this molecular epidemiology investigation, no identifying specimen data or clinical details were collected. Sex, date of birth and date of specimen collection were retained in a database. Patients could not be reidentified from this database. In response to a clinical request, specimen RNA extracts were provided by Pathology Queensland Central laboratory and were screened for *Parechovirus A* at the Qpid laboratory. RNA was extracted from 200 µl of sample using the Total Nucleic Acid Kit in the MagNAPure (Roche, Germany; eluted in 100 µl) or the DX buffer kit in the X-tractorGene (Qiagen, Australia; extract eluted in 130 µl) according to workflow and batching.

### Parechovirus Adetection

RNA extracts were screened using a previously described RT-rtPCR which targets the 5’UTR.[27] The screening HPeV RT-rtPCR employed 100 nM of each oligonucleotide primer (AN345_panHPeV/LV, AN344_panHPeV/LV; Geneworks, Australia) and 100nM of a FAM-BHQ1 dual-labelled fluorogenic oligoprobe (AN257_HPeV/LV; Geneworks, Australia) in a 20 µl one-step RT-PCR reaction mix (SensiFAST OneStep Mix; Bioline, Australia) including RNase inhibitor and 5 mM MgCl_2_. After 2 µl of nucleic acid extract was added and reverse transcribed for 20 min at 45°C, the mixes were incubated at 94°C for 2 min then cycled through 45 rounds of 94°C for 15 s, 58°C for 30 s and 72°C for 10 s using a RotorGene 3000, 6000 or RGQ (QIAGEN, Australia) to acquire fluorescence data at the final (annealing) step of each cycle.

### Parechovirus A genotvping

HPeV-positive specimens were amplified using a conventional nested RT-PCR assay incorporating previously described primers that bracket an approximately 256nt stretch, encompassing part of the viral protein coding regions VP3 and VP1 between nucleotide 2,182 and 2437 (numbering excludes primers and is based on prototype sequence L02971).[24] In our experience, this region is more likely to be successfully amplified than the VP1 region. Each experiment included a no-template (water) control and an HPeV-1 positive control (previous positive sample extract). The RT-PCR included 600 nM external sense and antisense oligonucleotide primers (HPeV_VP3/1_OS, HPeV_VP3/1_OAS, Geneworks, Australia) in a 20 µl one-step RT-PCR reaction mix (SensiFAST OneStep Mix, Bioline, Australia) with RNase inhibitor and 3 mM MgCl_2_. Two microliters of RT-PCR products were added to 18µl PCR reaction mixes (MyTaq HS DNA Polymerase; Bioline, Australia) containing 380 nM internal sense and antisense oligonucleotide primers (HPeV_VP3/1_IS, HPeV_VP3/1_IAS) and 4.75 mM MgCl2. Mixes were incubated at 94°C for 1 min then cycled through 40 rounds of 94°C for 30 s, 50°C for 30 s and 72°C for 105 s followed by a 7 min incubation at 72°C. Amplicons were electrophoresed to confirm presence and size of amplicon then sequenced (BigDye sequencing kit v3.1, Applied Biosystems Pty. Ltd; Australian Equine Genetics Research Centre, The University of Queensland). Following *in silico*removal of the primer sequence (Geneious Pro v8.1)^14^, *Parechovirus A* sequences were assigned GenBank accession numbers KX579839-KX579861.

Sequences were compared to the GenBank database by BLASTn analysis and to other *Parechovirus A* sequences after multiple alignment and phylogenetic analysis. A sequence was assigned to an HPeV genotype after meeting two criteria. Firstly, the new sequence had to cluster with a picornavirus study group (PSG)-assigned defined genotype in a phylogenetic analysis. Secondly, the new sequence had to share 90% or greater nucleotide sequence identity with the relevant region of a fully sequenced PSG-characterised genotype.

### Approvals

This work was completed under the approvals issued by Queensland Children’s Health Services human research ethics committee (#HREC/10/QRCH/95) and University of Queensland Medical Research Ethics Committee (#2010001381).

## Results

### Screening results

From 62 extracts submitted to our laboratory with a clinical suspicion of an HPeV infection between November 2013 and April 2014, 25 were *Parechovirus A* positive by RT-rtPCR. Of these 25, 23 extracts from 22 patients could be genotyped. These 23 extracts originated from CSF (n=14; 61% of all genotypes), faeces/stool (n=6; 26%), throat swab (n=2; 9%) or nasopharyngeal aspirate (NPA; n=1; 4%) samples. Most samples were received during the first month of summer 2013 (December 2013; n=22), a month which yielded 55% of all HPeV positive samples (n=12; Figure 1A). The average age of the patients infected with a defined HPeV genotype was 29 days (range 2 days to 90 days; Figure 1B).

**Figure 1.**
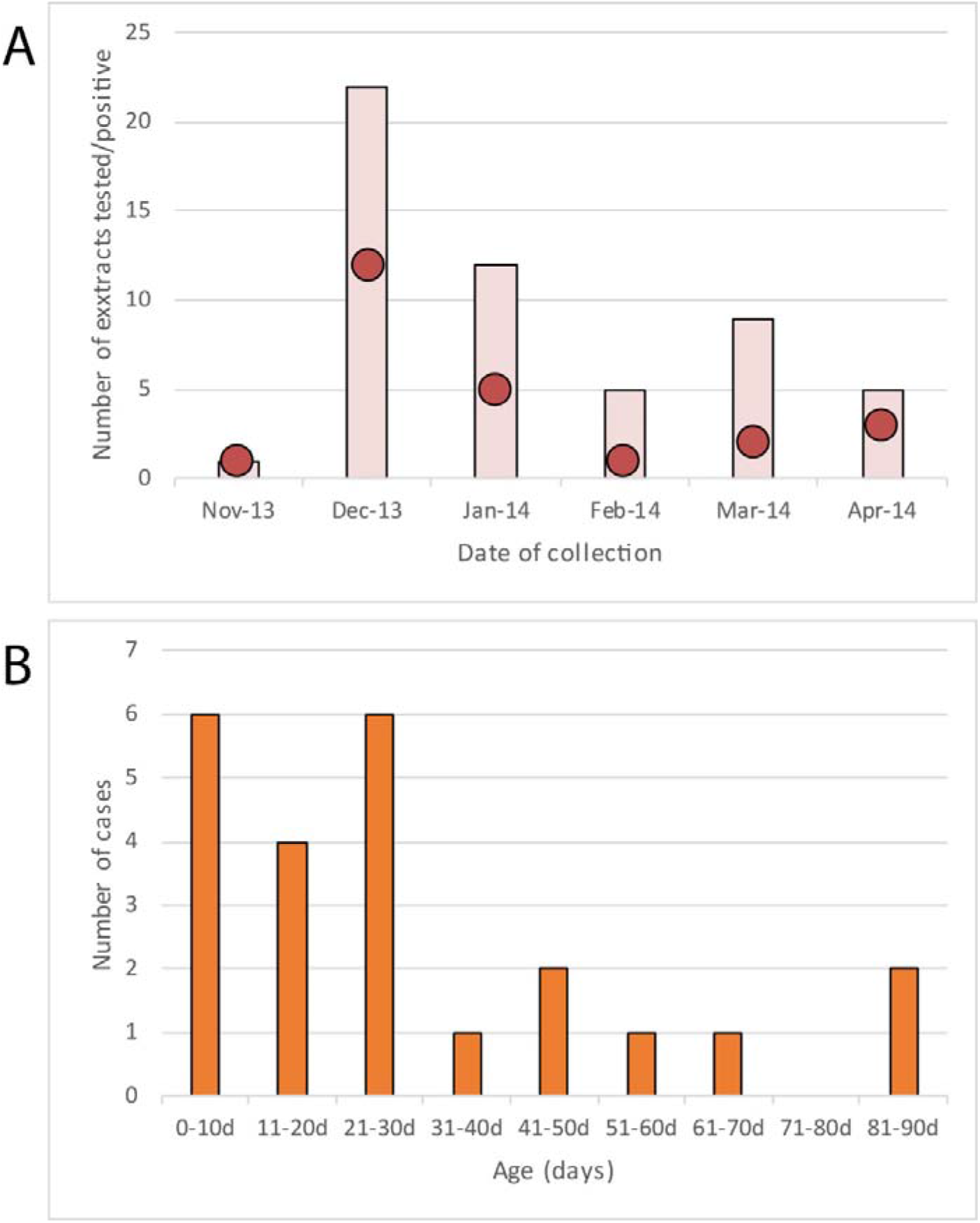
Epidemiology of HPeV detections. A. Month when testing was requested (bar) and the number of HPeV positive samples by month (red circle). B. Age distribution of the HPeV-positive samples.

### HPeV genotype identification

To identify the HPeV genotype, phylogenetic analysis, comparison of nucleotide sequences and of percent nucleotide identities were examined using sequences from published HPeV variants of previously assigned HPeV types.[24] Some HPeV genotypes have only VP1 sequence (required for genotype assignment) publicly available and few or no complete genomes or VP3/VP1 subgenomic sequences. Therefore, the reliability of the VP3/VP1 region for genotyping of all *Parechovirus A* genotypes remains untested. As noted elsewhere,[28] the HPeV1 genotype exists as two clades – lA and lB – also visible using the VP3/VP1 genotyping primer set of Simmonds et al.[24] The HPeV1 prototype Harris/USA-1956 (Genbank accession L02971) virus was initially the sole occupant of the lA branch, however, it has since been found that these viruses continue to circulate.[29-31] Interestingly, we found that the more recently described HPeV-17 genotype is also divergent; it’s VP3/VP1 sequences are interspersed among HPeV-4, 7 and 19 sequences(Fig 2). Both phylogeny and sequence identity matching using the VP3/VP1 regions requires comparison to characterised HPeV genotype sequences.

**Figure 2.**
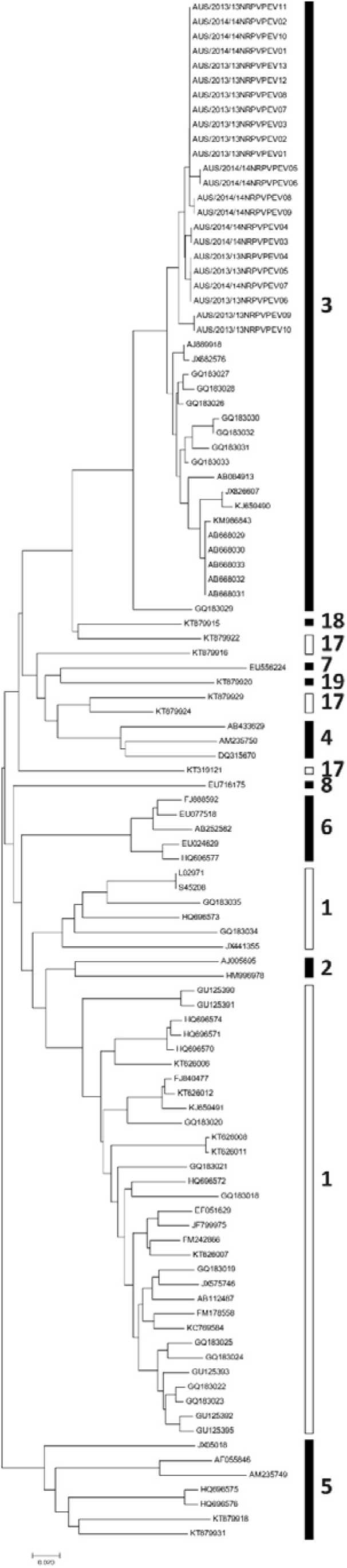
Phylogenetic analysis of the HPeV VP3/VP1 sequences (approximately 256nt in length) inferred using the Neighbor-Joining method [32] The percentage of replicate trees in which the associated taxa clustered together in the bootstrap test (500 replicates) are shown next to the branches [33] The evolutionary distances are presented as the number of base substitutions per site. All ambiguous positions were removed for each sequence pair. Open boxes indicate divergent genotypes with sequences intersected by those of other genotypes. Trees and evolutionary analyses were conducted using MEGA7 from alignments constructed using Geneious v8.1.8 and [34, 35].

A new guide tree of complete coding nucleotide sequences was prepared. (Supp Fig 1) The tree reflected current PSG HPeV numbering, confirming that the subgenomic VP3/VP1 region is not always accurate for phylogenetic representation of HPeV.

In addition to phylogenetic analysis, ≥90% nucleotide sequence identity between any new VP3/VP1 sequence and a characterised HPeV genotype sequence in the same region was used to support its genotype assignment.

All the HPeV positive samples characterised by this study were HPeV-3 variants, most closely matching those previously reported in Japan [36], Denmark [37], Italy [38], Scotland [9], Taiwan [39], England [26] and since described as a novel recombinant virus reported in Victoria, Australia.[40, 41] Whether the variants we reported here are also part of this recombinant virus cannot be confirmed from the phylogenetic analysis and BLAST comparisons of the VP3/VP1 subgenomic sequences we describe. However, it is noteworthy that they share the closest nucleotide identity (99%) of any HPeV-3 variants on GenBank.

Our observation that HPeV-3 was the predominant genotype in Queensland supports findings from a comprehensive study of samples collected over a 5-year period in Scotland.[9] We had previously described the detection of two HPeV instances in Brisbane from the NPAs collected in 2003 from ill children under two years of age.[42] We have since identified 0.5-1.0% of respiratory tract extracts from ill children in 2008-2012 inclusively, retrospectively tested positive for HPeV using the same assay described here (manuscript in preparation).

## Discussion

All of the *Parechovirus A* positive samples we sequenced contained variants of a recently described HPeV-3 genotype identified in infants during the spring and summer of 2013-14 in eastern Australia.

Our study had several limitations. Samples were not systematically collected or tested for other pathogens; they were received and tested upon the request of a medical officer and they were collected from ill patients. No estimate can be made of which other HPeV genotypes may have been missed but were circulating, or the frequency of mild and asymptomatic HPeV infection in Brisbane. While HPeV-3 is believed to cause infection and inflammation of the CNS, it is also detected in the gut of those with acute gastrointestinal disease and from the upper respiratory tract, where it has been associated with relatively mild respiratory illness.

Droplets and surface contamination may be a route of transmission, and while respiratory or gastrointestinal disease is not always present in PCR-positive children, it may be that the respiratory tract and oral cavity are viral landing sites from which dissemination of the infection may result in CNS infection/inflammation in some proportion of those infected.

HPeV molecular testing should be considered part of a systematic diagnostic panel to screen ill neonates and infants, especially when there are CNS-related symptoms present and during warmer months. The importance of this has recently increased with findings of serious sequelae in some proportion of HPeV cases.[13] Consideration could be given to assays that can specifically identify genotypes of most concern, however genotyping will not impact on supportive therapy or patient management and there is no antiviral or vaccine currently available for any HPeV. While recombination among the HPeVs means the subgenomic region used here is not ideal for genotyping in isolation, when combined with phylogenetic and BLAST analyses of characterized genomes, the VP3/VP1 region yields robust genotype identification. Knowledge of the pathogen may favor management of inflammation which may in turn result in fewer sequelae. Any future vaccine strategy will need to consider the early age of virus acquisition and severe disease attributed to HPeV-3.

Among young children presenting to hospital who present with suspected CNS infection/inflammation, seizures and/or rash, and where other pathogens have been ruled out, HPeV is an important consideration in the differential diagnosis.[10] The authors suggest that sensitive and specific routine surveillance for HPeVs be considered. The inclusion of HPeV screening could reduce the need for invasive sampling, complex testing, use of broad spectrum antibiotics and may speed patient discharge.

## Acknowledgements

We thank Pathology Queensland Central for the provision of specimen nucleic acid extracts and Dr Sarah Tozer, for provision of some data. This work was carried out within the Qpid laboratory, CHRC.

## Author Contributions

DM, CYTW and IMM conducted screening and genotyping. IMM designed the analysis and supervised DM and CYTW. IMM, KEA, AF and DM co-wrote the manuscript. All authors have read and accepted the manuscript.

## Supplemental data

**Supp Figure 1.**
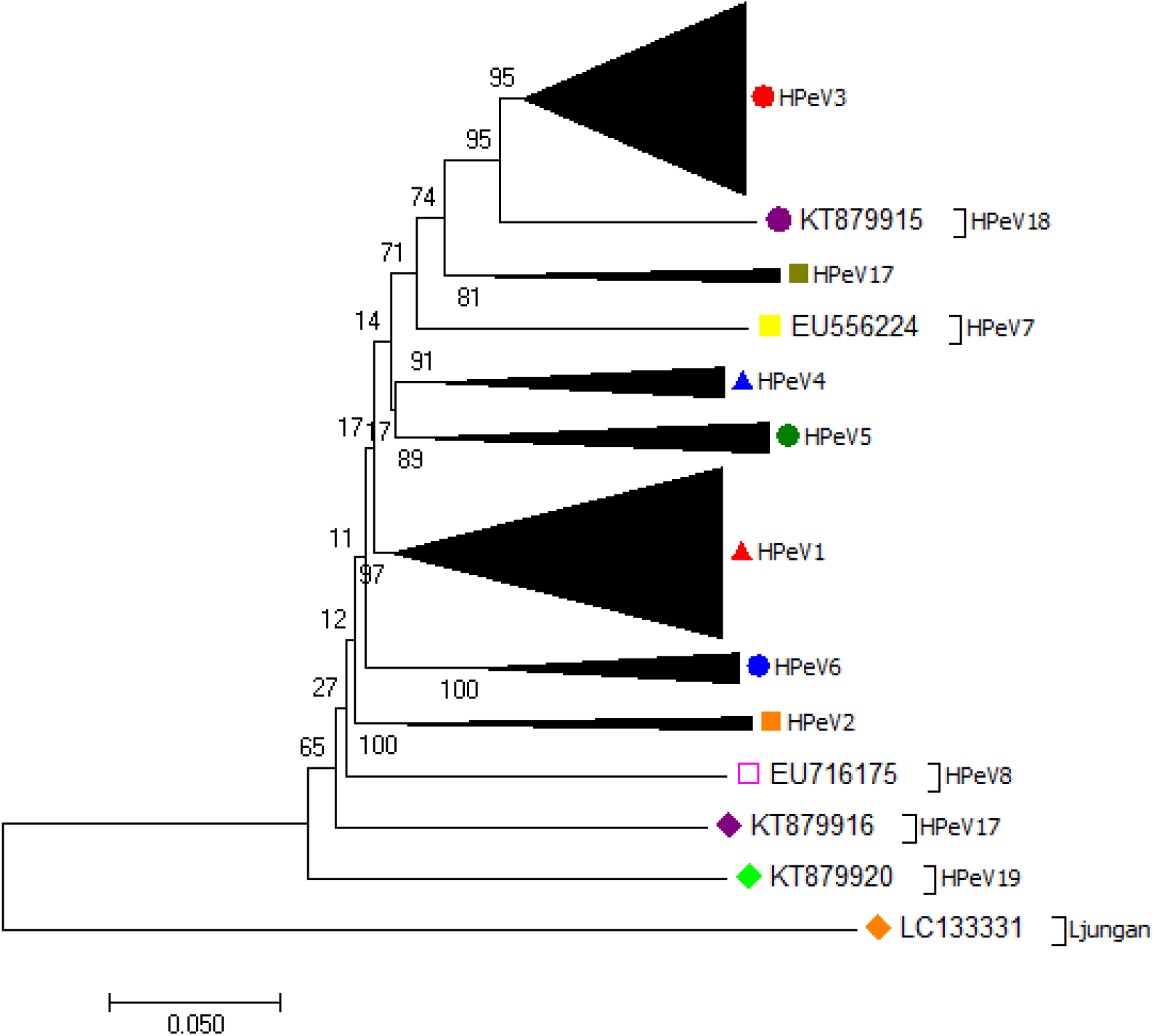
Phylogenetic relationships between available complete coding region nucleotide sequences.

The evolutionary history was inferred using the Neighbor-Joining method [1]. The percentage of replicate trees in which the associated taxa clustered together in the bootstrap test (500 replicates) are shown next to the branches [2]. The tree is drawn to scale, with branch lengths in the same units as those of the evolutionary distances used to infer the phylogenetic tree. The evolutionary distances were computed using the p-distance method [3] and are in the units of the number of base differences per site. The analysis involved 118 nucleotide sequences. All ambiguous positions were removed for each sequence pair. Evolutionary analyses were conducted in MEGA7 [4].

